# Characterization of *Plasmodium falciparum* myosin B in complex with the calmodulin-like domain of its light chain MLC-B

**DOI:** 10.1101/2022.05.28.493848

**Authors:** Isa Pires, Yu-Fu Hung, Ulrich Bergmann, Justin E. Molloy, Inari Kursula

## Abstract

Myosin B (MyoB) is a class 14 myosin expressed in all invasive stages of the malaria parasite, *Plasmodium falciparum*. It is not associated with the glideosome complex that drives motility and invasion of host cells. During red blood cell invasion, it remains at the apical tip of the merozoite but is no longer observed once invasion is completed. MyoB is not essential for parasite survival but, when it is knocked out, merozoites are delayed in the initial stages of red blood cell invasion, giving rise to a growth defect that correlates with reduced invasion success. Here, we have expressed and purified functional MyoB with the help of parasite-specific chaperones Hsp90 and Unc45, characterized its binding to actin and its known light chain MLC-B using biochemical and biophysical methods, and determined its low-resolution structure in solution using small-angle X-ray scattering. In addition to MLC-B, four other putative regulatory light chains were found to bind to the MyoB IQ2 motif *in vitro*. The purified recombinant MyoB adopted the overall shape of a myosin, exhibited actin-activated ATPase activity, and moved actin filaments *in vitro*. The ADP release rate was faster than the ATP turnover number, and thus, does not appear to be rate-limiting. This, together with the observed high affinity to actin and the specific localization of MyoB, may point towards a role in tethering and/or force sensing during early stages of invasion.

## Introduction

Malaria is an infectious disease caused by parasites of the genus *Plasmodium* that caused the death of more than 627 000 people in 2020 (1). Parasite motility, host cell invasion, and egress from the host cells is mediated by an actomyosin motor (2–4). Myosins are a diverse family of motor proteins composed of a heavy chain (100 to 150 kDa molecular weight) and a variable number of light chains (17-35 kDa). The heavy chain has three functional regions: A canonical N-terminal motor domain that is responsible for actin binding, ATP hydrolysis, and force generation; a light-chain-binding region (lever arm) with IQ motifs where one or more calmodulin-like light chain subunits bind; and a highly variable C-terminal tail region with sites that determine the specific activities of a particular myosin. The tail region may have coiled-coil forming motifs that lead to heavy chain oligomerization and filament formation and/or globular domains that specify cargo binding, regulation of motor activity, and subcellular targeting (5–7). Genomic analysis of the *Plasmodium* spp. identified six myosin genes (5), of which myosin A (MyoA) and myosin B (MyoB) are both class 14 myosins that drastically differ from other myosin classes in that they lack the tail region (2, 7, 8). MyoA is the best studied as it is present in the glideosome motor complex and is the main motor responsible for generating force for motility and invasion of the red blood cell (2, 9, 10). Two light chains have been identified for MyoA: A, so-called, essential light chain (ELC), which occupies the first IQ motif (IQ1), and MyoA tail domain interacting protein (MTIP), which occupies the second IQ motif (IQ2) (9, 11–14).

MyoB, the smallest *Plasmodium* myosin, with 17 fewer residues than MyoA (7) is expressed in all invasive stages (merozoites, ookinetes, and sporozoites) (8, 15). It is a stationary myosin, first found with a fixed localization in the apical portion of merozoites (7). Later, it was shown to localize more specifically in the extreme apex of merozoites, ookinetes, and sporozoites (8, 15), overlapping with tubulin, indicative of association with the polar rings (8). This is also supported by a ring-like shape in mature ookinetes, as seen in super-resolution microscopy (15). During invasion, MyoB remains at the apical tip of merozoites and disappears once the process is completed (8). MyoB does not colocalize with MyoA, and there is no evidence of association with the glideosome protein complex (15), even though *in silico* predictions (16) and subsequent interaction studies (17) have suggested the possibility of such interactions. MyoB is not essential for parasite survival (8, 15), but it has been hypothesized to be involved in erythrocyte invasion (7). MyoB conditional knockout merozoites showed reduced invasion success due to delayed initiation of parasite internalization, however, invasion into the red blood cell still proceeded, and viable parasites, albeit with a small growth defect, were produced (8, 10, 15).

A single light chain, MLC-B, has been identified for MyoB (8). In ookinetes, MLC-B is positioned at the apical end early during development, 12 h before MyoB is first detected (15). For a myosin light chain, MLC-B is large, being composed of an EF-hand-containing calmodulin-like domain at the C-terminus and a large N-terminal domain of unknown function, predicted to form a coiled-coil structure (8). Interestingly, unlike MyoB, MLC-B is essential for parasite survival, which together with its early appearance at the apical end might suggest additional functions, independent of MyoB (8, 15). It is also possible that another myosin could take over some functions and interact with MLC-B in place of MyoB.

In this study, we have produced recombinant *Plasmodium falciparum* MyoB in complex with MLC-B calmodulin-like domain (CaMLD) and performed biochemical and structural characterization of this protein complex. We show that MyoB can be expressed and purified in sufficient amount and purity for structural and biochemical studies, using a heterologous expression system, including parasite-specific chaperones for correct protein folding. The purified recombinant MyoB is an active motor with a higher affinity to actin than MyoA. The MyoB neck IQ2 region can interact, *in vitro*, with four calmodulin family light chains other than MLC-B.

## Results

### Parasite-specific chaperones are required for correct folding of MyoB

The expression of *P. falciparum* MyoB alone in *Spodoptera frugiperda* (Sf)21 insect cells resulted in very little or no soluble protein, despite extensive screening for solubilizing conditions (Fig. 1A). Co-infecting the cells with equal amounts of viruses for MyoB and the chaperone Unc45, as described previously for MyoA (14), or infecting with a virus containing MyoB, Unc45, and Hsp90, did not improve the solubility of the protein (Fig. 1A). Soluble MyoB could be obtained by infecting the cells with two viruses, one containing MyoB-Unc45-Hsp90 and another containing Unc45, at a ratio of 10:1 (v/v) (Fig. 1A). The overall expression level of MyoB decreased with the addition of chaperones (Fig. 1A), and although the amount of soluble protein on the gels does not seem to increase substantially with the addition of surplus Unc45, this was the only condition that enabled producing useful amounts of soluble MyoB.

**Figure 1.**
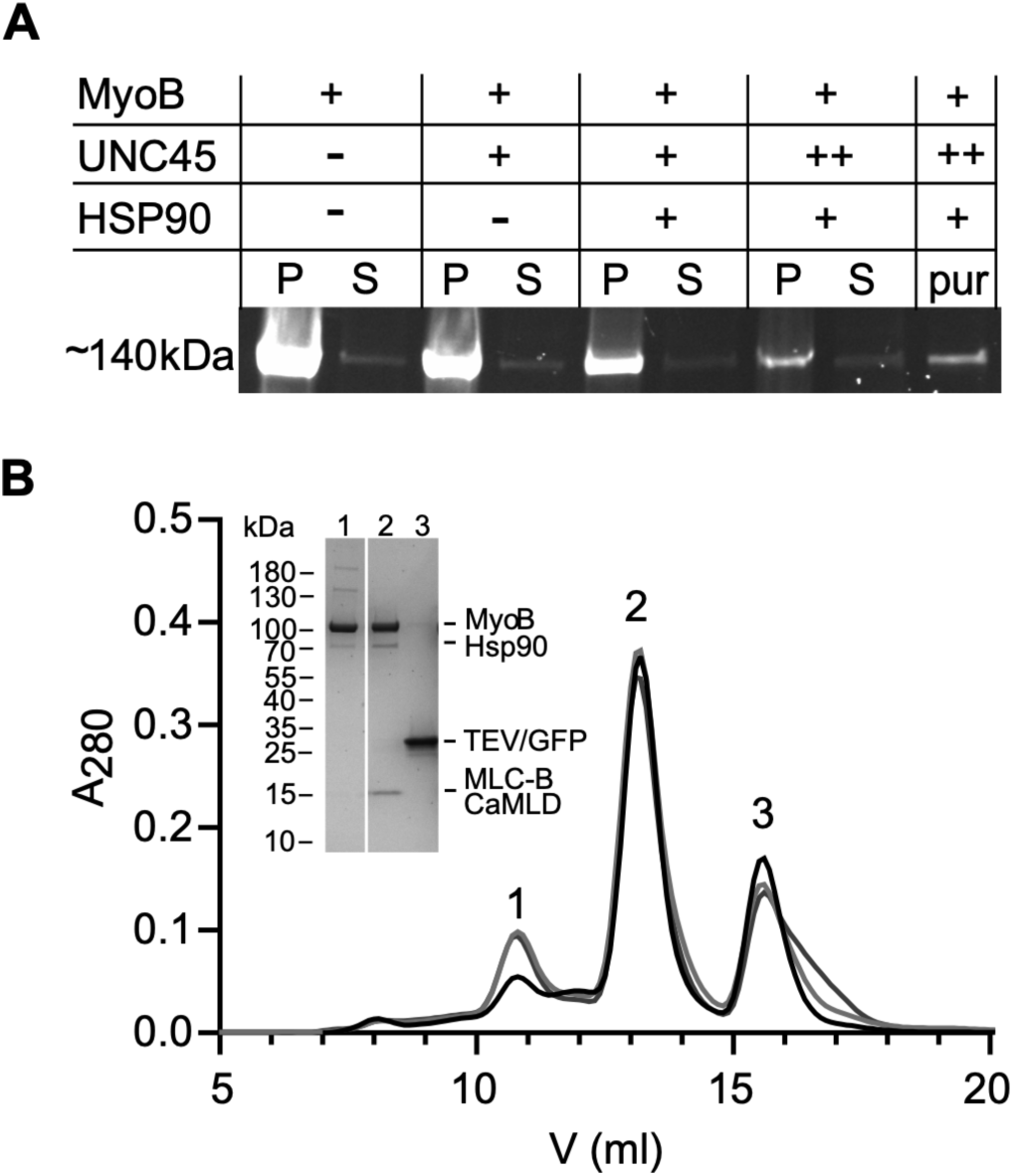
MyoB solubility and the MyoB-MLCB CaMLD complex purification. A, SDS-PAGE imaged under UV light for detection of eGFP-MyoB fusion protein. S: soluble fraction; P: insoluble fraction; pur: purified protein. Equal culture and buffer volumes were used for all samples, with the exception of pur that acts as a control. B, Elution of MyoB-MLC-B CaMLD from a Superdex S200 10/300 column in 20 mM Tris-HCl (pH 8.5), 150 mM KCl, 0.5 mM TCEP, 5 mM MgCl2, 0.1 mM EGTA at a flow rate of 0.35 ml min^-1^. The different black-grey curves represent three independent purifications. The inset shows samples from the SEC peaks on SDS-PAGE. Peak 1 contains partially unfolded MyoB, peak 2 the folded MyoB-MLC-B CaMLD complex, and peak 3 Strep-tag-eGFP/TEV.

We and others (8) have been unable to purify full-length MLC-B. Likewise, copurification of this light chain with MyoB failed. Negatively stained gold-particle immune electron micrographs of the *Sf*21 expression cells revealed that full-length MLC-B is severely aggregated *in cellulo* (Fig. S1). Therefore, we used for all experiments a construct where the N-terminal coiled-coil region of the full-length MLC-B was removed, thus resulting in just the C-terminal CaMLD (hereafter termed MLC-B CaMLD) (8).

The stability of the neck region of myosins is often increased by the binding of light chains (18). Here, MyoB and MLC-B CaMLD were produced in different expression systems. Therefore, purified MLC-B CaMLD was added to the MyoB cell lysate immediately after cell disruption by sonication and before clarification by centrifugation. Even though the lysis buffer had a high ionic strength (500 mM NaCl), MLC-B CaMLD still bound to MyoB (Fig. 1B). Size exclusion chromatography (SEC) showed that the fraction of MyoB not bound to MLC-B CaMLD eluted earlier than the MyoB-MLC-B CaMLD complex, together with Unc45 and Hsp90 (Fig. 1B). This suggests that this small amount of MyoB may be partially unfolded and/or aggregated (Fig. 1B). The MyoB-MLC-B CaMLD complex eluted as a symmetric peak (Fig. 1B).

### MyoB-MLC-B CaMLD complex is folded and has an overall shape of a myosin

Synchrotron radiation circular dichroism spectroscopy (SR-CD) was used to evaluate whether the purified proteins were folded and to determine their thermal stability. Deconvolution of SR-CD spectra indicated that the MyoB-MLC-B CaMLD complex purified as above is composed of 36% α-helix, 14% β-sheet, 13% turn, and 37% other structure, as calculated using the BeStSel server (19). The complex is therefore folded (Fig. 2A), with secondary structure contents similar to other myosins (20). The normalized root-mean-square deviation was low (0.006), indicating a strong correlation between the data and the calculated secondary structure contents. The melting temperature (T_m_) of the complex was 50.6°C, calculated from a sigmoidal fit to the peaks at 223 and 194 nm (Fig. 2B).

**Figure 2.**
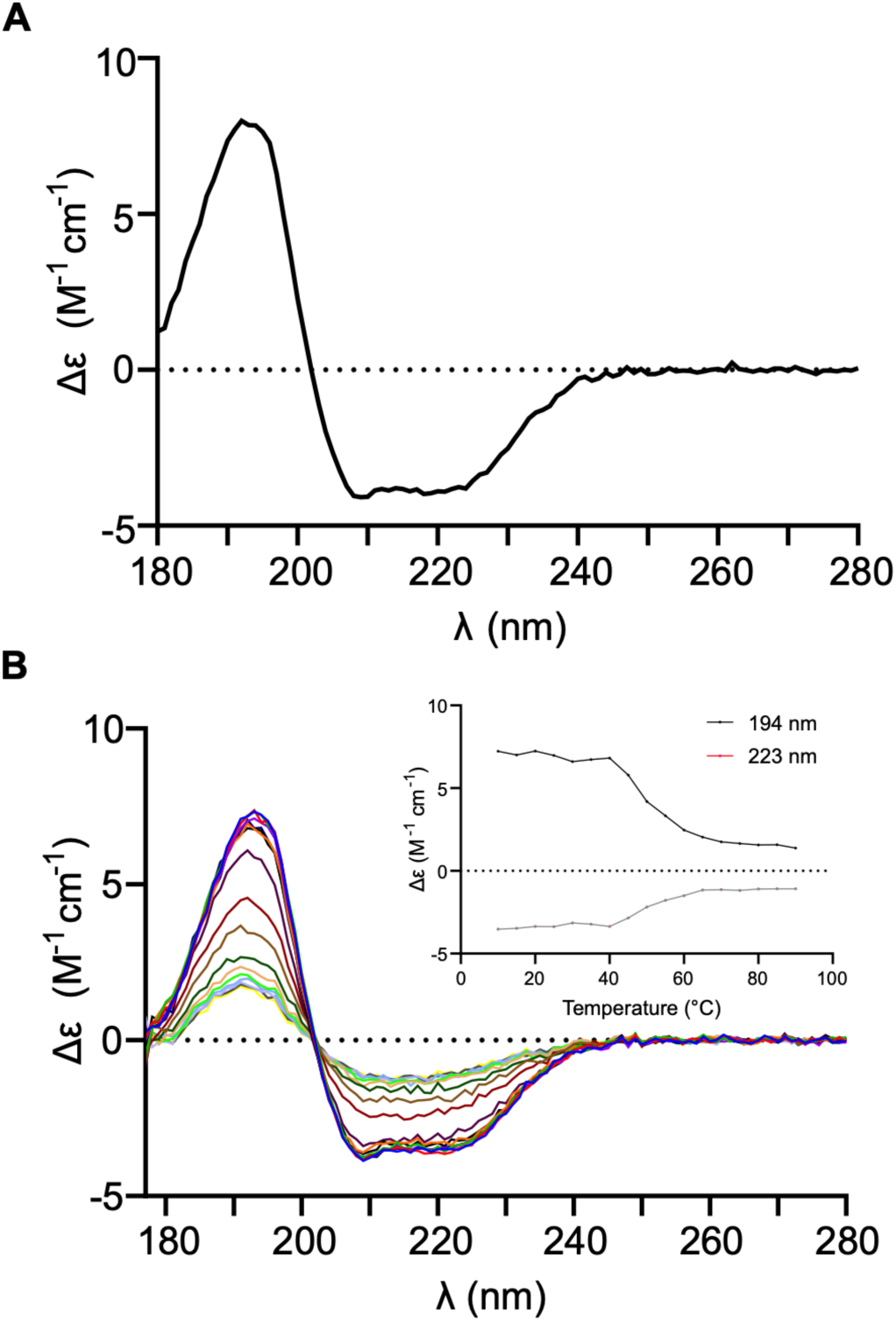
MyoB-MLC-B complex folding and thermal stability. A, SR-CD spectrum of the MyoB-MLC-B CaMLD complex. B, SR-CD melting spectra of MyoB-MLC-B CaMLD complex. The inset represents the T_m_ of MyoB-MLC-B CaMLD from calculation of a sigmoidal fit to the peaks at 223 and 194 nm.

We used small-angle X-ray scattering (SAXS) to determine the shape and dimensions of the MyoB-MLC-B CaMLD complex in solution (Fig. 3). The shape of the scattering curve and the dimensionless Kratky plot indicate a folded, elongated protein with some flexible parts (Fig. 3A,B). The observed maximum distance (D_max_) of over 200 & Å shows that the complex is rather extended (Fig. 3C, Table S1). Low-resolution *ab initio* models generated using DAMMIN (21) and GASBOR (22) show an overall shape that fits well to a monomeric myosin motor domain with a bound light chain (Fig. 3D). A recent high-resolution structure of the MyoA light-chain complex (23) also matches the shape and dimensions of the SAXS models (Fig. 3D).

**Figure 3.**
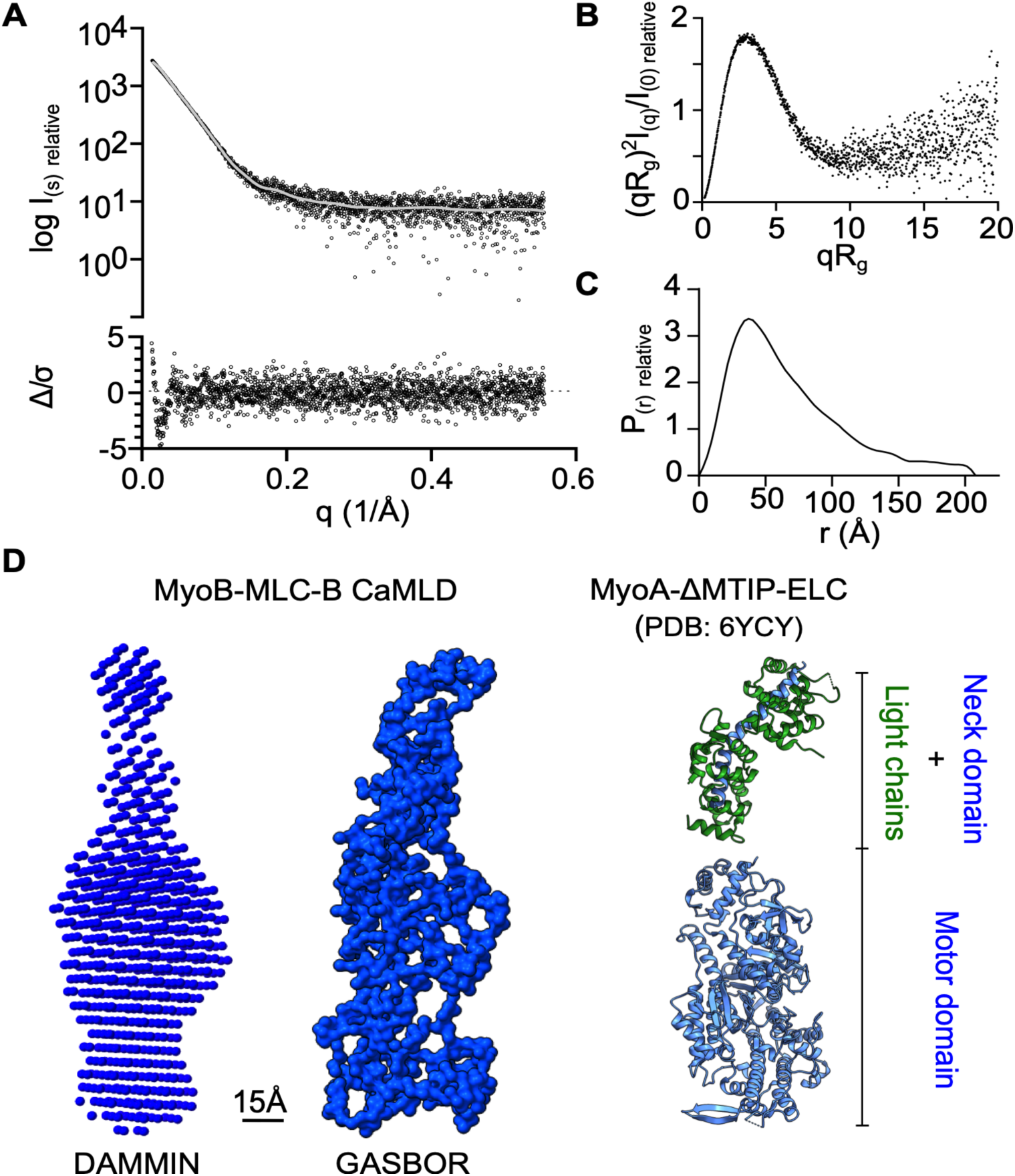
SAXS of the MyoB-MLCB CaMLD complex. A, SAXS data (black dots, upper panel) of MyoB-MLCB CaMLD. The fit of the GASBOR model to the experimental data is shown as a grey line in the upper panel. The residuals are denoted in the lower panel, where Δ/σ= [I_exp_(q)- I_mod_(q)]/σ(q). B, The dimensionless Kratky plot suggests a folded, elongated shape of the complex C, Distance distribution function of the MyoB-MLC-B CaMLD complex. D, Comparison of *ab initio* models of the MyoB-MLC-B CaMLD complex from DAMMIN (left) and GASBOR (middle) to the crystal structures of MyoA neck-MTIP-ELC (upper right; PDB: 6YCY; Moussaoui et al., 2020) and MyoA motor domain (lower right; PDB: 6YCY; Moussaoui et al., 2020). Note that the neck region with the light chains is shown in an approximate extended conformation, unlike the compact form seen in the crystal structure, likely affected by crystal packing.

*In vivo*, MyoA can be phosphorylated at Ser19, and this post-translational modification (PTM) is required for maximum motility (24). The interaction between Ser19 and Lys769 stabilizes the rigor-like state of MyoA, modulating the force and the speed at which it moves along actin (24). Sequence alignment shows that MyoB has a serine at position 16 (close to Ser19 of MyoA) while Lys769 in MyoA corresponds to Arg747 in MyoB (Fig. S2). The molecular weight of recombinant MyoB was determined by mass spectrometry to be 93376 Da, which corresponds very closely to the expected size 93378.86 Da without any PTMs. Thus, Ser16 does not appear to be phosphorylated in the recombinantly produced MyoB.

### MyoB interacts with MTIP and three other putative light chains

*In silico* predictions and subsequent experimental data have suggested that the MyoA regulatory light chain MTIP can bind to MyoB (16, 17). However, there have been no *in vitro* experimental data on the second MyoA light chain *Pf*ELC binding to MyoB. Here, pulldowns of MTIP and ELC using N-terminally Strep-tagged MyoB neck (Ala742-Glu801) were performed. While there was no evidence of ELC binding to the MyoB IQ motifs (Fig. 4A, B), MTIP was pulled down using MyoB (Fig. 4A, B), as reported previously (17). It is worth noting that after coexpression and following the normal purification protocol for MyoB-MLC-B CaMLD (500 mM NaCl lysis), MTIP did not copurify with MyoB, suggesting that this interaction is weaker than that with MLC-B CaMLD. Although it was possible to pull down MTIP with MyoB *in vitro*, there is no evidence of this interaction taking place *in vivo*.

**Figure 4.**
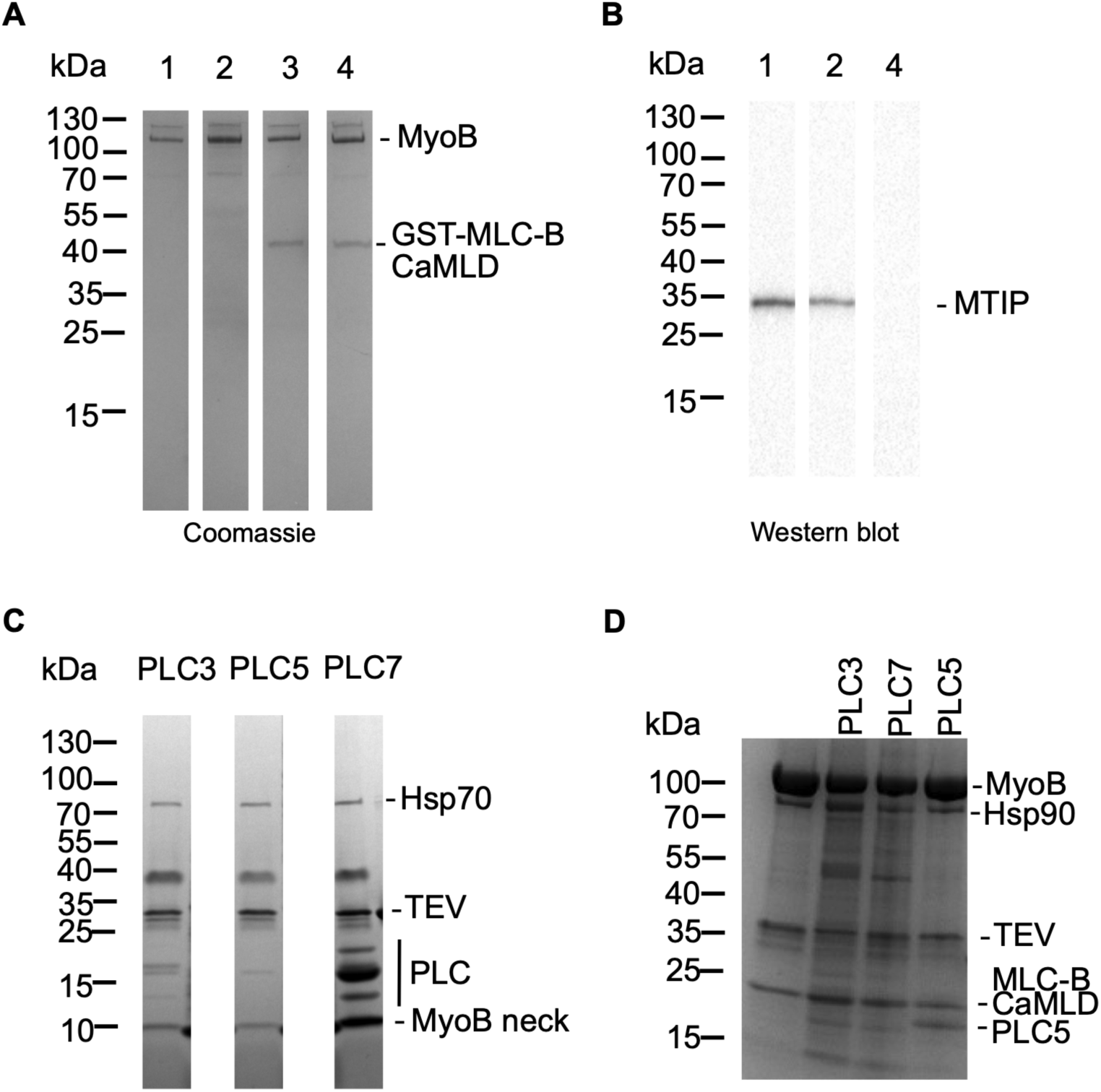
ELC identification. A, Coomassie brilliant blue stained SDS-PAGE gel showing the binding of MTIP, ELC, and MLC-B CaMLD to MyoB. 1: MyoB-MTIP; 2: MyoB-MTIP-ELC; 3: MyoB-GST-MLC-B CaMLD; 4: MyoB-MLC-B CaMLD-ELC B, Western blot performed using the same samples as the Coomassie brilliant blue stained gel shown on panel A. Anti-His antibody was used to verify the presence or absence of MTIP and ELC. C, Binding of PLCs to MyoB neck domain. PLC3, PLC5 and PLC7 were identified as light chains interacting with MyoB. D, Pulldown of PLC3, PLC5 and PLC7 using the MyoB-MLC-B CaMLD complex. The bands not labeled in the figure are contaminants from insect cell expression system.

For MyoA, MTIP binding is necessary for ELC to interact (23, 25). Hence, pulldown of ELC with the MyoB-MLC-B CaMLD complex was also attempted using the same strategy as above. However, there was still no evidence of ELC binding to the available IQ motif of the MyoB neck (Fig. 4A, B). It seems likely that ELC cannot bind to MyoB in the presence or absence of MTIP/MLC-B CaMLD.

The *P. falciparum* genome encodes several putative light chains (PLC). Two approaches were used to identify PLCs binding to MyoB. First, for faster screening, insect cells were coinfected with viruses containing the MyoB neck and one PLC at a time, to assess if any of the PLCs bound to either of the possibly available IQs. Three PLCs, namely PLC3 (PF3D7_0414200), PLC5 (PF3D7_0627200), and PLC7 (PF3D7_1434200), were able to bind to the MyoB neck (Fig. 4C), and their identities were confirmed by mass spectrometry. Knowing that IQ2 is occupied by MLC-B in the parasite, pulldowns of PLC3, PLC5, and PLC7 separately using a pre-formed purified complex of MyoB-MLC-B CaMLD were performed (Fig. 4D). Only PLC5 was able to bind to the pre-formed MyoB-MLC-B CaMLD complex. PLC3 and PLC7 did not bind in those conditions. Thus, it is probable that their interaction with MyoB in the absence of MLC-B CaMLD occurs through IQ2.

As only PLC5 could be pulled down with the MyoB-MLC-B CaMLD complex, we attempted to purify the ternary complex by incubating the purified MyoB-MLC-B CaMLD complex with PLC5 and an excess of MLC-B CaMLD, followed by concentration and SEC. The SEC elution profile exhibited two peaks in the presence or absence of Ca^2+^, and PLC5 was not associated with the MyoB-MLC-B CaMLD complex (Fig. S3). Thus, the successful pulldowns of PLC5 by MyoB (Fig. 4C, D) were most probably due to PLC5 interacting with IQ2 and competing with MLC-B CaMLD (Fig. 4C, D). Also, the binding of either light chain is not dependent on Ca^2+^. In conclusion, we have not been able to identify any light chain binding to the MyoB IQ1 region but several light chains can bind to its IQ2 *in vitro*.

### MyoB-MLC-B CaMLD binds actin and moves actin filaments *in vitro*

Consistent with our biochemical and biophysical analyses, negatively stained electron microscopy (EM) images of MyoB-MLC-B CaMLD indicated that the protein complex was homogeneous and monomeric, with no evidence of Y-shaped, two-headed, myosin dimers (Fig. 5A). Jasplakinolide-stabilized *P. falciparum* actin I filaments mixed, in the absence of Mg.ATP, at a ratio of 5:1 with the MyoB-MLC-B CaMLD complex appeared more rugged than jasplakinolide-stabilized actin I filaments alone and showed punctate localization of myosin-like structures on the filaments (Fig. 5A). However, fully decorated, arrowhead-like, structures as seen for MyoA (26) were not observed. Ultra-centrifugation sedimentation assays, performed in the absence of Mg.ATP, showed that MyoB bound to and co-sedimented with porcine skeletal muscle α-actin (Fig. 5B). We conducted *in vitro* motility assays using fluorescence video microscopy to characterize the ability of MyoB to move actin filaments in the presence of Mg.ATP. MyoB was captured on a microscope coverslip either by binding nonspecifically to a surface layer of nitrocellulose or by using an anti-GST antibody to capture the GST-tagged MLC-B CaMLD. Both methods allow the myosin head domain to face upwards. The antibody-capture method was performed in two different ways: 1) The microscope coverslip with surface-immobilized anti-GST antibody was first incubated with MLC-B CaMLD and subsequently incubated with MyoB. 2) The antibody-coated coverslip was incubated directly with the fully formed MyoB-MLC-B CaMLD complex. In the absence of Mg.ATP, immobile, rhodamine-phalloidin labelled porcine α-actin filaments were observed tightly bound to the coverslip surface. Upon addition of 2 mM Mg.ATP, the actin filaments moved in a directed manner, and a small number of non-moving filaments were also observed. The different antibody capture methods resulted in no difference in actin filament gliding speed (0.23 μm s^−1^ for method 1; 0.21 μm s^−1^ for method 2). Thus, the data sets were combined for further analysis. When MyoB-MLC-B CaMLD was non-specifically immobilized on a nitrocellulose-coated surface, the actin filament gliding speed was 0.12 ± 0.53 μm (N_obs_ = 4301), which is about two-fold slower than the antibody-capture speed 0.22 ± 0.02 μm s^−1^ (N_obs_ = 3965) (Fig. 5C, Movie 1, Movie 2). MyoB in the absence of its MLC-B CaMLD light chain immobilized to a nitrocellulose coated surface bound actin tightly in the absence of Mg.ATP but showed no filament gliding upon ATP addition (Movie 3). Showing that, as expected, the light chain is necessary for the stabilization of the so-called “lever arm” region and production of force and movement (27). The speed observed for MyoB-MLC-B CaMLD was similar to that of MyoA with a single MTIP light chain (14) but about 10-fold slower compared to other studies with slightly different experimental conditions (9, 23, 28).

**Figure 5.**
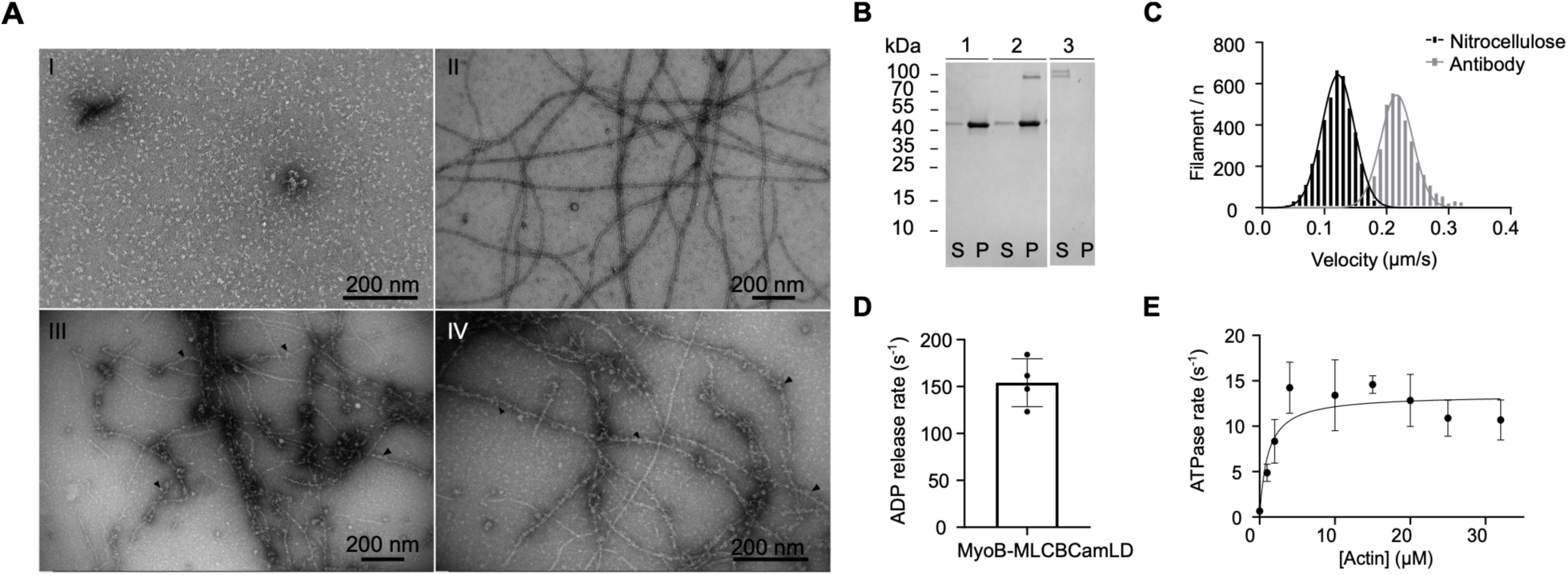
MyoB binding to actin. A, I -Negative-stained electron micrograph of MyoB-MLC-B CaMLD at 100 nM. II-Negative-stained electron micrograph of JAS-stabilized *P. falciparum* actin I at 500 nM. III, and IV - Negative-stained electron micrographs of MyoB-MLC-B CaMLD binding to filamentous ActI, at a ratio of 5:1. Scale bars: 200 nm. Arrowheads indicate possible MyoB-MLC-B CaMLD particles. B, Cosedimentation of MyoB with F-α-actin. The double band of MyoB is due to the lack of bound MLC-B which allows for a degradation of the neck region. 1: F-α-actin control; 2: actomyosin; 3: MyoB control; S: supernatant; P: pellet. C, Motility assay of the MyoB-MLC-B CaMLD complex using nitrocellulose and antibody capture. Nitrocellulose captured data are colored black and the antibody captured data are colored grey. The antibody-capture method exhibited faster motility. D, The ADP release rate of the MyoB-MLC-B CaMLD complex was 154 ± 25 s^-1^, at 24°C (four independent experiments). E, Actin-activated steady-state ATPase activity of the MyoB-MLC-B CaMLD complex showed 20-fold lower K_app_, but similar k_cat_ values to the MyoA-MTIP complex (four independent experiments). All assays were performed using filamentous α-actin.

The rate of dissociation of the tightly-bound acto-MyoB-MLC-B CaMLD rigor complex, following ATP addition, was determined by incubating α-actin labeled with N-(1-pyrene)iodoacetamide (NPI), MyoB-MLC-B CaMLD, and ADP and then rapidly mixing in Mg.ATP at 24°C using a stopped-flow apparatus. The dissociation rate of MyoB-MLC-B CaMLD from α-actin, as measured by the fluorescence increase of NPI, which reports the rate of ADP release from the motor protein, was 154 ± 25 s^-1^ (Fig. 5D). This is slightly slower than the previously obtained rate for MyoA at 25°C (209 ± 39 s^-1^) (24). In a separate experiment, the steady-state actin-activated ATPase activity of the MyoB-MLC-B CaMLD complex was measured (Table 1). The “apparent” K_m_ (K_app_) for the actin-activated ATPase activity was 1.4 μM, and the maximum actin-activated ATPase activity (k_cat_) was 13 s^−1^ (Fig. 5E). The k_cat_ was similar to the value obtained for MyoA at 23°C (14) and for fast skeletal muscle myosin 2 (29), but the K_app_ was approximately 20-fold lower than for MyoA (9, 14, 24), indicating a higher affinity to actin. Similarly, the so-called coupling efficiency (k_cat_/K_app_) was ∼9, which is at least 2 orders of magnitude higher than reported for most myosins (Table 1) (14, 24, 29–35). The rate of ADP release from MyoB (154 s^-1^) was ∼10-fold faster than the maximum actin-activated ATPase activity, indicating that ADP release is unlikely to be the rate-limiting step in the MyoB ATPase cycle.

**Table 1:**
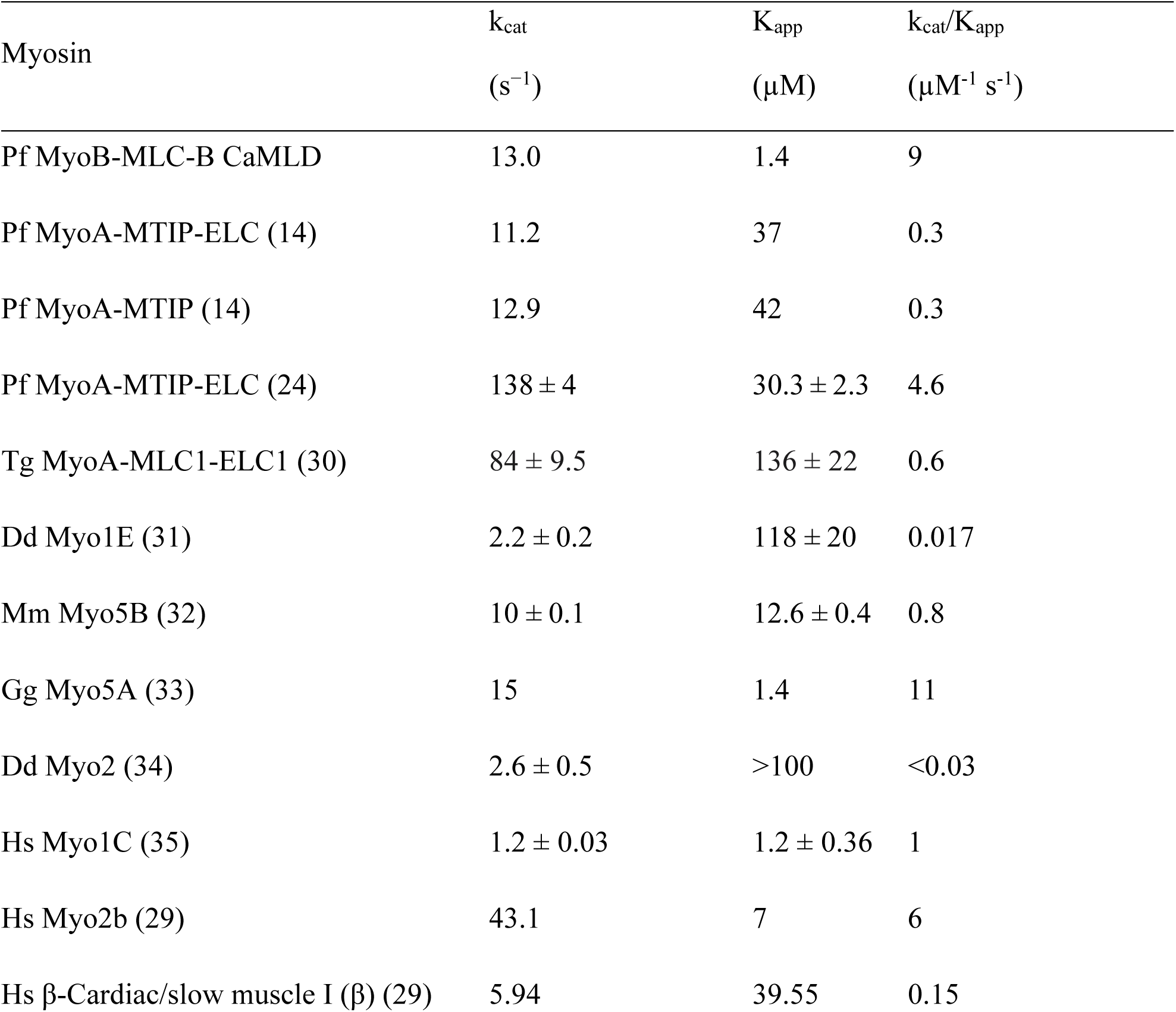
Steady-state kinetic parameters of MyoB-MLC-B CaMLD in comparison with selected previously characterized myosins.

| Myosin | $k_{\text{cat}}$<br>($\text{s}^{-1}$) | $K_{\text{app}}$<br>($\mu\text{M}$) | $k_{\text{cat}}/K_{\text{app}}$<br>($\mu\text{M}^{-1} \text{s}^{-1}$) |
| --- | --- | --- | --- |
| Pf MyoB-MLC-B CaMLD | 13.0 | 1.4 | 9 |
| Pf MyoA-MTIP-ELC (14) | 11.2 | 37 | 0.3 |
| Pf MyoA-MTIP (14) | 12.9 | 42 | 0.3 |
| Pf MyoA-MTIP-ELC (24) | $138 \pm 4$ | $30.3 \pm 2.3$ | 4.6 |
| Tg MyoA-MLC1-ELC1 (30) | $84 \pm 9.5$ | $136 \pm 22$ | 0.6 |
| Dd Myo1E (31) | $2.2 \pm 0.2$ | $118 \pm 20$ | 0.017 |
| Mm Myo5B (32) | $10 \pm 0.1$ | $12.6 \pm 0.4$ | 0.8 |
| Gg Myo5A (33) | 15 | 1.4 | 11 |
| Dd Myo2 (34) | $2.6 \pm 0.5$ | >100 | <0.03 |
| Hs Myo1C (35) | $1.2 \pm 0.03$ | $1.2 \pm 0.36$ | 1 |
| Hs Myo2b (29) | 43.1 | 7 | 6 |
| Hs β-Cardiac/slow muscle I ($\beta$) (29) | 5.94 | 39.55 | 0.15 |

## Discussion

This study aimed to shed light on the *P. falciparum* class 14 myosin, MyoB, and its interaction with several partner proteins. While much research is being conducted on MyoA, other *Plasmodium* myosins are less well studied, and their interaction partners and functions are to a large extent unknown.

### Chaperones and myosin light chains are essential for MyoB stability and function

Genetic analyses show that myosins are a diverse family of actin-based motor proteins found in all eukaryotic organisms. Myosins can be divided into ∼35 distinct structural classes (36). (Odronitz & Kollmar, 2007). The best-studied, class 2 myosins, were first discovered and isolated from smooth, skeletal, and cardiac muscle tissue. Our understanding of the diverse classes of myosins has been underpinned by ability to express and purify protein in heterologous expression systems. Access to recombinant protein permits biophysical and biochemical analysis of myosins that may be present in only minute quantities in the native organism. Some myosins can still only be produced in mammalian tissue-culture systems (*e*.*g*. class 2 striated muscle myosins), while others require co-expression of specific chaperones for correct folding in baculovirus/insect cell expression systems (30, 37). The Unc45-Hsp90 chaperone complex facilitates myosin motor domain folding; Hsp90 binds Unc45 through its tetratricopeptide repeat domain and Unc45 binds myosin through its C-terminal regions. Unc45 interacts with several types of myosins, and Hsp90 binds folding intermediates shortly before their native conformation is achieved. (38–40)

Two interesting findings in this work were that MyoB, when compared to MyoA, required a higher ionic strength buffer for isolation (9, 14, 23, 30) and two chaperones for proper folding (Fig. 1A). The fact that it was necessary to infect the cells with an additional copy of the Unc45 virus to obtain soluble, correctly folded, MyoB, suggests that MyoB folding requires a high local concentration of Unc45, possibly the formation of a scaffold of Unc45 proteins, offering multiple adjacent binding sites for heat-shock cognate protein 70, Hsp90, and myosin, as seen in *Caenorhabditis elegans* (41). As typical for myosins, also MyoB requires at least one light chain for full functionality. With a molecular weight of 78 kDa, MLC-B is an unusually large light chain, compared to other known myosin light chains, which are in the range of 17-35 kDa. As MyoB, like also MyoA, lacks the entire tail domain, it is a tempting hypothesis that the N-terminal predicted coiled-coil domain, for which no function has been assigned, could perform some functions that typically would be carried out by the myosin tail, such as oligomerization or targeting to membranes or certain cellular compartments. The MLC-B N-terminal domain is prone to aggregation both *in vitro* and *in cellulo* in the recombinant expression host insect cells. Given the punctate localization of MyoB *in vivo* in the parasites, this aggregation or formation of large agglomerates might be relevant for the anchoring of MyoB to the very apical tip of the parasite (8, 15).

The secondary structure contents of the MyoB-MLC-B CaMLD complex (Fig. 2, Table S2) were comparable to those in homologous crystal structures, such as that of myosin V (20) (PDB: 1W7I, chain A, Table S2). The T_m_ of the MyoB-MLC-B CaMLD complex was slightly higher (Fig. 2B, Table S3) compared to the T_m_ values of other myosins (Table S3). This could indicate that the nucleotide-free state of MyoB is more stable than the motor domains of the previously characterized myosins. However, the stability of MyoB was measured in the presence of its light chain, contrary to the other tested myosins, which might influence the T_m_ value.

In MyoA, phosphorylation of Ser19 and its interaction with Lys769 (Fig. S2) have an impact on the speed and force produced by the motor (24). Ser16 in MyoB aligns with Ser19 of MyoA (Fig. S2B), but Lys769 in MyoA is replaced by Asp748 in MyoB in the sequence alignment and the predicted structure of MyoB (Fig. S2A). This substitution reverses the charge, which would have implications on the equivalent interaction in MyoB. Our mass spectrometry results indicate that the MyoB heavy chain has no PTMs and that the protein complex characterized here is unphosphorylated. It should be noted that Ser16 may be phosphorylated *in vivo*, but it seems likely that its possible effect on the myosin function may not be the same or may not follow the same mechanism as in MyoA.

The *ab initio* SAXS model of MyoB-MLC-B CaMLD shows the classical shape of a myosin motor domain with some additional volume that could correspond to MLC-B CaMLD (Fig. 3D). The dimensions of the model are close to those of the MyoA-MTIP-ELC crystal structure (23) if the neck with the bound light chains were in an extended conformation, not seen in the crystal structure, due to crystal packing. Additional flexibility due to the lack of a light chain bound to the IQ1 of MyoB may contribute to our failure to crystallize full-length MyoB with or without MLC-B CaMLD. The crystal structure of MyoA was obtained in complex with both light chains (23) or as a motor-domain only construct, without the neck region (24).

### MyoB-MLC-B CaMLD is an active motor

The MyoB-MLC-B CaMLD complex binds to actin and moves actin filaments in an *in vitro* gliding assay. The gliding velocity depended upon the method of surface immobilization, with specific antibody attachment to the CaMLD light chain GST tag giving the fastest speed. Previous studies using *Pf*MyoA have shown that the maximal gliding speed depends upon the method of surface attachment, heavy chain phosphorylation state, light chain complement and temperature (9, 14, 24, 28). Our current results with MyoB-MLC-B CaMLD were obtained using similar conditions to our previous work with MyoA (14). We find that the maximum actin gliding speed obtained using MyoB-MLC-B CaMLD is similar to that of MyoA-MTIP. It can be extrapolated that with the addition of an ELC to MyoB, the speed might double, as was seen for MyoA (9, 14, 24). However, we have not been able to identify an ELC for MyoB, and it is not clear whether one exists.

The maximum actin-activated ATPase activity determined for MyoB-MLC-B CaMLD, is similar to values found for MyoA (14) and similar to fast skeletal muscle myosin II (29). MyoB shows a 20-fold lower K_app_ and at least 2 orders of magnitude higher k_cat_/K_app_ for actin-activation indicating stronger affinity to actin. This may have biological importance, given that MyoB appears to be a stationary myosin at the apical tip, involved in internalization of the parasite during invasion (10). The higher affinity to actin is possibly due to sequence differences in the so-called hypertrophic cardiomyopathy (HCM) loop, in particular the TEDS site, where instead of a Thr, which is an important phosphorylation site in many myosins, is a Gln in MyoB. Interestingly, this residue is a Gln in many of the apicomplexan myosins. In fact, *P. falciparum* MyoA, which has a Thr in that position, seems to be an exception within the phylum. Of the two *P. falciparum* MyoA-actin I cryo-EM structures available, one is of the wild-type protein (26) and in the other one, the TEDS site Thr has been mutated to Asp, to mimic phosphorylation (42). In the wild-type structure, this side of the HCM loop, including Thr417, is not directly involved in contacts with actin. In the mutant, the Oδ atom of the mutated Asp is within 4.7 and 4.2 Å from Lys337 and Tyr338 of actin, respectively, and could be involved in a hydrogen bond or salt bridge interaction with either of the side chains. A Gln at this site, as in MyoB and many other apicomplexan myosis, does not provide a negative charge (like phosphorylation or Asp/Glu) that would enable salt bridge interactions but could form hydrogen bonds to both Lys337 and Tyr338, thus contributing to the actomyosin interface. The ADP dissociation rate from MyoB on actin is approximately 10 times faster than the actin-activated steady-state ATPase rate, implying that P_i_ release or some other step in the acto-myosin ATPase cycle is rate-limiting.

Taken together, our data and the cellular localization of MyoB suggest that it may serve an anchoring function during parasite invasion, helping to form and release the initial parasite-host cell contacts at the early stages of the moving junction formation. This would be analogous to the better-studied Myosin 1B, which is a low-duty-ratio, force-sensitive anchor, which enables control of membrane tension and deformation (43). MyoB knock-out parasites take longer to invade (10), indicating that the myosin substituting for MyoB is not an effective motor for that purpose.

### Several light chains bind to the MyoB neck region

*In vitro*, MTIP can bind to MyoB (Fig. 4B) as also reported before (17), but we could not detect any binding for *Pf*ELC (Fig. 4A-B). The IQ2 (Fig. S2) of MyoA has two important residues, Gln808 and Arg812, involved in MTIP binding (44), and those residues are conserved in MyoB. On the other hand, there are several residues essential for *Pf*ELC binding to MyoA IQ1 (Fig. S2) that are not conserved in MyoB IQ1. Cys780 of MyoA, which interacts with the C-lobe of ELC (23), is replaced by Leu760 in MyoB. Val781 of MyoA, which interacts with the intra-lobe linker of ELC (23), is a cysteine in MyoB. Thus, it is not clear whether there is an ELC that can bind to IQ1 of MyoB and whether such a light chain would be needed for maximum motility, like in MyoA.

In a simple pulldown experiment, we identified four additional light chains, which can bind to MyoB IQ2 (Fig. 4C, D). Only further *in vivo* colocalization studies will be able to clarify the biological significance of this discovery. It can be hypothesized that the identified putative light chains may be used in the schizont or sporozoite forms, where MyoB also showed cytoplasmic distribution, besides the apical ring localization (15), or that they can be used to stabilize the myosin neck until it is bound to MLC-B. Indeed, the expression information available for these light chains indicate expression in schizonts and for PLC7 also in sporozoites (45, 46). Importantly, we have not yet been able to identify a specific light chain that binds to IQ1 of MyoB, so it remains unclear whether this IQ motif is occupied. However, we believe it is highly unlikely that this region of the MyoB heavy chain would not have an associated light chain *in vivo*.

### Concluding remarks

One of the biggest obstacles to studying *Plasmodium* myosins has been in producing correctly-folded, functional material using traditional methods of recombinant protein production. Here, we show that it is necessary to optimize the ratio of chaperones to myosin to produce sufficient amounts of *P. falciparum* MyoB for biochemical and biophysical analysis. This may be important for producing other myosins, as well. We have determined the actin activation of MyoB ATPase activity as well as its *in vitro* gliding motility, ADP release, and ATP hydrolysis rates. Our results are directly comparable to our previous work with MyoA and show that the affinity of MyoB to actin was markedly stronger than that of MyoA. This, together with the fact that the ADP dissociation rate from MyoB on actin is ten times faster than the actin-activated steady-state ATPase rate, implies that P_i_ release or another step preceding ADP release is rate-limiting and supports a tethering/force sensing function *in vivo*. To clarify this would require more detailed kinetic and functional, including *in vivo* analyses. Finally, we found that IQ2 of MyoB was also able to bind several putative *Plasmodium* myosin light chains, but *in vivo* studies are needed to find out their biological relevance.

## Experimental procedures

### Protein expression and purification

*P. falciparum* MyoB (*PF*3D7_0503600), chaperones Unc45 (*PF*3D7_1420200) and Hsp90 (*PF*3D7_0708400), PLC3 (PF3D7_0414200), PLC5 (PF3D7_0627200) and PLC7 (PF3D7_1434200), codon-optimized for *Sf*21, were cloned by sequence and ligation independent cloning or the restriction enzyme method into a baculovirus transfer vector (pFL, Fitzgerald et al., 2006). The codon-optimized *Hsp90* was a kind gift from Dr. Will Stanley (University of Edinburgh). Strep-tag (IBA GmbH) and eGFP were used as N-terminal tags for MyoB. MyoB was co-expressed with Unc45 and Hsp90 to achieve correct folding, as suggested previously (30). The construct was introduced to Max Efficiency DH10Bac cells (Invitrogen), bacmids were isolated, and the gene integrations were confirmed using standard protocols (Bac-to-Bac, Invitrogen). *Sf2*1 cells (Gibco) were transfected using FuGene 6 (Promega). The supernatant containing V_0_ virus (virus passage zero (progeny virus)) was collected by centrifugation after 7 days of incubation at 27°C and 325 rpm. Working virus passage 1 stocks were prepared by infecting *Sf*21 suspension cell cultures with V_0_ and harvesting the supernatants after 4-5 days of incubation at 27°C and 100 rpm. Protein expression was started by infecting *Sf*21 suspension cell cultures at densities between 1.0 and 2.0×10^6^ cells ml^-1^. For MyoB expression, a ratio of 10:1 (v:v) of MyoB-Unc45-Hsp90:Unc45 viruses was used. The culture was harvested at 72 h post-infection. *Sf*21 cells expressing MyoB were lysed by mild sonication using a tapped disruptor horn (Branson 450 Digital Sonifier), in 20 mM Tris-HCl pH 8.5, 500 mM NaCl, 0.5 mM tris(2-carboxyethyl)phosphine (TCEP), 10 mM MgCl_2_, 1 mM EGTA, 1× SigmaFast EDTA-free protease inhibitor, followed by the addition of previously purified MLC-B CaMLD. The lysate was clarified by centrifugation at 38 000 g for 45 min and then allowed to flow-through Streptactin XT matrix (IBA) twice, at 4°C. One wash with lysis buffer without protease inhibitors was followed by two consecutive washes with 20 mM Tris-HCl pH 8.5, 250 mM NaCl, 0.5 mM TCEP, 5 mM MgCl_2_, 1 mM EGTA and 20 mM Tris-HCl pH 8.5, 150 mM NaCl, 0.5 mM TCEP, 5 mM MgCl_2_, 1 mM EGTA. The protein was eluted by on-column cleavage of the tag with tobacco etch virus protease (TEV) and subsequently concentrated using a molecular weight concentrator (Vivaspin). The protein was filtered with a 0.22 μm pore size filter (Millipore) prior to size-exclusion chromatography (SEC) that was carried out in a Superdex S200 10/300 GL column (GE Healthcare), equilibrated with SEC buffer (20 mM Tris-HCl pH 8.5, 150 mM KCl, 0.5 mM TCEP, 5 mM MgCl_2_, 0.1 mM EGTA). If not used immediately, the concentrated elution fractions were flash-frozen using liquid N_2_ and stored at -70°C.

The MLC-B CamLD plasmid was kindly provided by Drs. Judith Green and Anthony Holder, and the purification followed as previously described (8). Protein was purified and stored pure at -20°C with or without GST-tag. MTIP (*PF*3D7_1246400) and ELC (*PF*3D7_1017500) were expressed and purified as described elsewhere (Green et al., 2017). *P. falciparum* actin I and skeletal muscle α-actin were expressed and purified as previously described (Kumpula et al., 2017).

### Mass spectrometry

Protein masses were determined using liquid chromatography–mass spectrometry with an Aquity UPLC system (Waters) linked to a Q-ExactviePlus mass spectrometer. A BioResolveRPmABPolyphenyl column 2.1×50 mm was developed with a gradient comprising 0.1% Formic acid (A) and 0.5% Formic acid in acetonitrile (B) with a flow rate of 0.4 ml/min and initial conditions 5% B; 0.5 min: 10% B; 9 min: 60% B; 10min 80% B; 11min 5% B. The mass spectrometer was operated in positive mode with resolution 17500 and m/z range 400 to 4000. Automatic Gain Control and maximum injection time were set to 3×10^6^ and 200 msec, respectively. Raw data were processed with BioPharmaFinder (Thermo), using the Respect option with standard settings suitable for proteins of MyoB molecular weight.

### Synchrotron radiation circular dichroism spectroscopy

The secondary structure contents and folding state of the MyoB-MLC-B CaMLD complex were assessed using SR-CD. Protein concentrations between 1.0-1.7 mg ml^-1^, previously dialyzed into a far-UV compatible buffer (10 mM Tris-HCl pH 8.5, 150 mM KF, 5 mM MgSO_4_, 0.5 mM TCEP) were used. CD spectra were recorded between 170-280 nm at the AU-CD beamline on ASTRID2 at ISA (Aarhus, Denmark) at 10°C in 100-μm pathlength closed circular cells. Thermal stability spectra were obtained from 10-90°C, using a 5°C increment. Data were deconvoluted using the BeStSel server (19), using smoothed data between 175-250 nm. T_m_ data were calculated from a sigmoidal fit to the peaks at 223 and 194 nm using GraphPad Prism 9.

### Small-angle X-ray scattering

SAXS was used to analyze low-resolution structural features of MyoB-MLC-B CaMLD. SAXS data were collected at the P12 BioSAXS beamline at the PETRA III storage ring (EMBL/DESY, Hamburg, Germany). A concentrated sample of MyoB-MLC-B CaMLD at 3 mg ml^-1^ was dialyzed against the buffer, 20 mM Tris-HCl pH 8.5, 150 mM KCl, 5 mM MgCl_2_, 0.1 mM EGTA, 0.5 mM TCEP, for 6 h at 4°C. A dilution series and their corresponding buffers were measured at room temperature. The automatically processed data were further analyzed using ScÅtter (47) and the ATSAS package (48). *Ab initio* models were generated using DAMMIN (Svergun, 1999) and GASBOR (22). ChimeraX 1.1 was used for preparing the figures (49).

### Pulldown assays

For pulldown assays, MyoB, MLC-B CaMLD and the known MyoA light chains, MTIP and ELC, were purified as described previously, without removing the tags. The proteins were incubated with each other in different combinations at a concentration of 20 μM for 30 min at room temperature, as follows: MyoB-MTIP-ELC; MyoB-MTIP, MyoB-MLC-B CaMLD and MyoB-MLC-B CaMLD-ELC. MyoB and any interacting light chains were bound to Streptactin XT resin *via* the MyoB Strep-tag, in 20 mM Tris-HCl pH 8.5, 150 mM KCl, 5 mM MgCl_2_, 0.1 mM EGTA, 0.5 mM TCEP. The resin was washed with 50 column volumes of the same buffer. Elution was performed by the addition of 50 mM biotin to the same buffer. Samples were taken of all steps and prepared for SDS-PAGE and Western blotting analysis.

For the putative light chain initial screening, the MyoB neck domain (Ala742-Glu801) was coexpressed with the different PLCs in *Sf*21 cells. The PLCs contained a His-tag with a human rhinovirus 3C protease cutting site, and MyoB neck domain contained a Strep-tag with a TEV cutting site. The cell pellets were resuspended in lysis buffer (20 mM Tris-HCl pH 7.5, 250 mM NaCl, 0.5 mM TCEP, 5 mM imidazole, 1 mM CaCl_2_, 1× SigmaFast EDTA-free protease inhibitor) and lysed by sonication (Branson 450 Digital Sonifier). The lysate was clarified by centrifugation (38 000 g, 45 min). The soluble fraction was incubated with equilibrated with HisPur nitrilotriacetate nickel agarose resin resin (Thermo) for 1 h at 4°C, under gentle agitation. One wash with lysis buffer without protease inhibitors was followed by two consecutive washes 20 mM Tris-HCl pH 7.5, 500 mM NaCl, 0.5 mM TCEP, 10 mM imidazole, 1 mM CaCl_2_ and 20 mM Tris-HCl pH 7.5, 150 mM NaCl, 0.5 mM TCEP, 15 mM imidazole, 1 mM CaCl_2_. The protein complexes were eluted by on-column cleavage of MyoB neck domain using TEV and subsequently concentrated using a centrifugal concentrator (Millipore). The samples were analyzed using SDS-PAGE, and gel bands representing PLCs were excised from the gel and analyzed using mass spectrometry. After the identification of the interacting PLC, full-length MyoB with or without MLC-B CaMLD bound was purified as described earlier until the wash step 2, then the purified PLC was allowed to incubate with the resin-containing MyoB for at least 30 min, followed by two consecutive washes with 20 mM Tris-HCl pH 7.5, 500 mM NaCl, 0.5 mM TCEP, 10 mM imidazole, 1 mM CaCl_2_ and 20 mM Tris-HCl pH 7.5, 150 mM NaCl, 0.5 mM TCEP, 15 mM imidazole, 1 mM CaCl_2_. On-column cleavage was used as an elution strategy using 20 mM Tris-HCl pH 7.5, 150 mM NaCl, 0.5 mM TCEP, 15 mM imidazole, 1 mM CaCl_2_ supplemented with TEV.

### Actin cosedimentation assays

To assess the binding of MyoB-MLC-B CaMLD to α-actin and *P. falciparum* actin I, a cosedimentation assay was performed. α-actin or actin I in G-buffer (10 mM HEPES pH 7.5, 0.2 mM CaCl_2_, 0.5 mM ATP, and 0.5 mM TCEP), in the absence or presence of MyoB, were polymerized overnight at room temperature, by adding F-buffer to final concentrations of 50 mM KCl, 4 mM MgCl2, and 1 mM EGTA, in addition to the G-buffer components, in a volume of 100 μl. Filaments were sedimented at 435 000 g for 1 h at room temperature. Both pellet and supernatant fractions were examined using SDS-PAGE with Coomassie brilliant blue staining.

### *In vitro* motility assays

*In vitro* motility assays were performed as previously described (14). With the following exceptions: the nitrocellulose-coated coverslips were incubated with protein or with anti-GST antibody (Invitrogen) for 4 min. The MyoB-MLC-B CaMLD complex was used at concentrations of 0.84 mg ml^-1^ for nitrocellulose capture, or 0.84 and 0.42 mg ml^-1^ for anti-GST tag capture.

### Actin-activated ATPase assay

The actin-activated ATPase of MyoB-MLC-B CaMLD was estimated using a nicotinamide adenine dinucleotide reduced (NADH)-coupled assay to quantify the rate of ATP hydrolysis, as described before (50). The assay buffer contained 100 units ml^-1^ lactate dehydrogenase, 500 units ml^-1^ pyruvate kinase, 2.5 mM phospho(enol)pyruvate, 2 mM ATP, and 0.3 mM NADH in KMg50 buffer (50 mM KCl, 2 mM MgCl2, 1 mM EGTA, 2 mM DTT, 10 mM imidazole, pH 7.0). The MyoB-MLC-B CaMLD complex was added to a final concentration of 0.15 μM, and jasplakinolide-stabilized filamentous α-actin was added at concentrations ranging between 0-25 μM. The rate of ATP hydrolysis was calculated from the rate of change in NADH absorbance, measured in a Tecan SPARK spectrophotometer at 340 nm. The experiments were conducted at 23°C. The data were fit to the Michaelis–Menten equation, using non-linear regression with a least-squares fit in GraphPad Prism 9.

### Transient kinetics

Transient kinetic experiments were performed in a stopped-flow spectrophotometer (Hi-Tech Scientific) to access the ADP release rate of MyoB. A monochromator with a 2 nm bandpass was used for fluorescence excitation, and a 400 nm cutoff filter was used to measure fluorescence emission. NPI α-actin, labeled as described before (51), was excited at 365 nm. All experiments were performed in 20 mM Tris-HCl pH 8.5, 150 mM KCl, 0.1mM EGTA, 5 mM MgCl_2_, 0.5 mM TCEP at 24 °C. The concentrations after mixing in the stopped-flow cell were 1 μM MyoB, 1 μM α-actin, 250 μM ADP, and 2 mM ATP. The ADP rates were obtained with the software provided with the instrument (Kinetic Studio 5), where 10-15 traces per protein preparation (4 independent protein preparations) were averaged before fitting to an exponential. Nonlinear least squares fitting of the data was conducted in Graphpad 9.0.

### Negative stain electron microscopy

The interaction of MyoB and filamentous α-actin was analyzed by diluting both proteins into 20 mM Tris-HCl pH 8.5, 5 mM MgCl_2_, 0.1 mM EGTA, 150 mM KCl, 0.5 mM TCEP. Filamentous α-actin (final concentration of 500 nM) was mixed with myosin (final concentration of 100 nM), and 5 μl of the sample were applied and incubated for 45 s on a glow-discharged formvar-coated copper grid (standard hexagonal grid with mesh size 300) with a layer of evaporated carbon. The formvar coating and carbon evaporation were carried out in-house. After incubation, the excess liquid was blotted with filter paper, and the grid stained twice with 1% (w/v) uranyl acetate (5 and 60 s, respectively), the excess liquid was blotted using filter paper, and the grid was left to dry in air. The grids were imaged using a Tecnai G2 Spirit microscope at 120 kV.

## Data availability

All the relevant data and plasmids used to support the findings of this study are available upon request from the corresponding authors.

## Supporting information

This article contains supporting information.

## Acknowledgments

We are thankful to Dr. Judith Green and Dr. Tony Holder for discussions and help with MLC-B and the PLCs, Dr. Andrea Lopez Moreno and Ju Xu for help with cloning the PLCs, and Dr. Petri Kursula for critical reading of the manuscript. We thank Dr. Arne Raasakka for collecting the SR-CD data, Dr. Juha Vahokoski for the collection of the motility assay images, and Dr. Ilkka Miinalainen, for helping with negative stain imaging at the Biocenter Oulu electron microscopy core facility. We also acknowledge the use of the AU-CD beam line on ASTRID2 at ISA (Aarhus, Denmark) for SR-CD measurements and the P12 BioSAXS beamline at the PETRA III storage ring (EMBL/DESY, Hamburg, Germany) for access to SAXS beam time.

## Author contributions

Isa Pires - Conceptualization, Formal analysis, Investigation, Methodology, Validation, Visualization, Writing – original draft

Yu-Fu Hung – Methodology, Supervision, Writing – review & editing Ulrich Bergmann – Methodology, Writing – review & editing

Justin Molloy – Formal analysis, Validation, Writing – review & editing

Inari Kursula - Conceptualization, Formal analysis, Funding acquisition, Project administration, Supervision, Visualization, Validation, Writing – review & editing

## Funding and additional information

This work was funded by the Academy of Finland (IK), Emil Aaltonen foundation (IK), Sigrid Jusélius foundation (IK), the Norwegian Research Council (IK) and Francis Crick Institute core funding (JEM); Cancer Research UK (FC001119), UK Medical Research Council (FC001119), Wellcome Trust (FC001119).

## Abbreviations and nomenclature

CaMLD: calmodulin like-domain
ELC: essential light chain
HCM: hypertrophic cardiomyopathy
Hsp: heat-shock protein
MLC-B: myosin light chain B
MTIP: myosin A-tail domain interacting protein Myo myosin
NPI: N-(1-pyrene)iodoacetamide
*Pf*: *Plasmodium falciparum*
PLC: putative light chain
PTM: post translational modifications
SAXS: small-angle X-ray scattering
SDS-PAGE: sodium dodecyl sulfate polyacrylamide gel electrophoresis
SEC: size-exclusion chromatography
*Sf*: *Spodoptera frugiperda*
SR-CD: synchrotron radiation circular dichroism spectroscopy
TCEP: tris(2-carboxyethyl)phosphine
TEV: tobacco etch virus protease Unc45 uncoordinated mutant number 45

