## Supplemental figures and tables for "Characterization of *Plasmodium falciparum* myosin B in complex with the calmodulin-like domain of its light chain MLC-B"

Material included:

Figure S1

Figure S2

Figure S3

Table S1

Table S2

Table S3

Legends of Movie 1, Movie 2 and Movie 3.

**Figures**

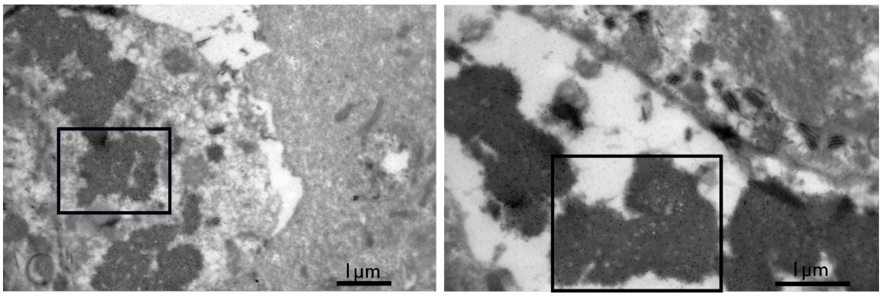


**Figure S1.** Immuno-negative staining electron microscopy micrographs of *Sf*21 cells at 48 h post-infection. Gold particles indicate the presence of His-tagged MLC-B, example highlighted with red box. Scale bar: 1 μm.

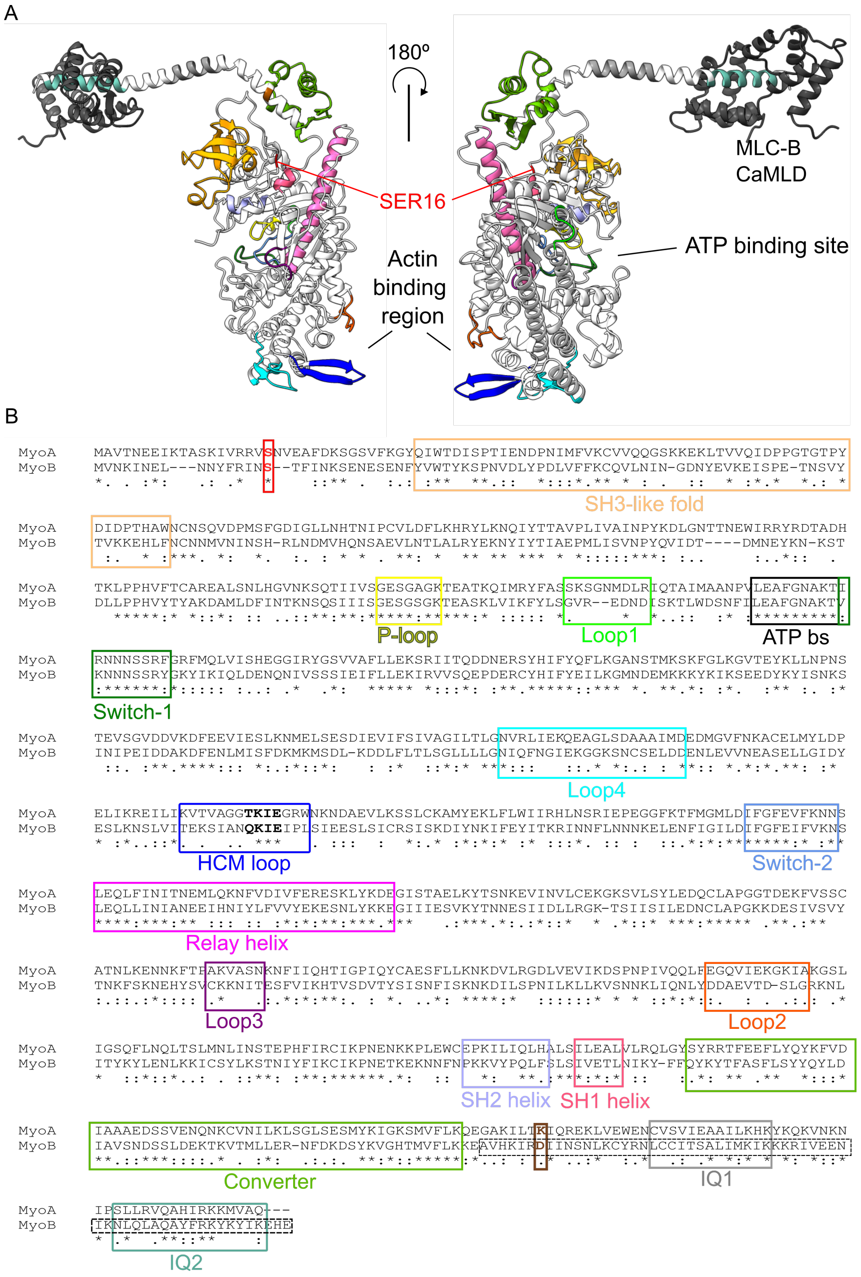
A
180º
SER16
MLC-B
CaMLD
ATP binding site
Actin
binding
region
B
MyoA
MyoB
SH3-like fold
P-loop
Loop1
ATP bs
Switch-1
Loop4
HCM loop
Switch-2
Relay helix
Loop3
Loop2
SH2 helix
SH1 helix
Converter
IQ1
IQ2

**Figure S2:** A, MyoB structure prediction using Alphafold (51) with functional regions of the motor and lever highlighted. The MLC-B CaMLD structure prediction using Alphafold (51) was superimposed using MTIP-MyoA interaction as guide. B, Sequence alignment of MyoA and MyoB using T-COFFEE (52) with functional regions of the motor and lever highlighted corresponding to Fig. S2A (2, 24, 53). The residues that influence the force and the speed at which the MyoA moves along actin are shown in red (Ser19) and brown (Lys769) (24). The dotted line box indicates the MyoB neck sequence used in this study. The residues in bold inside the HCM loop box (blue) indicate the TEDS site.

**Commented [IP1]:** added

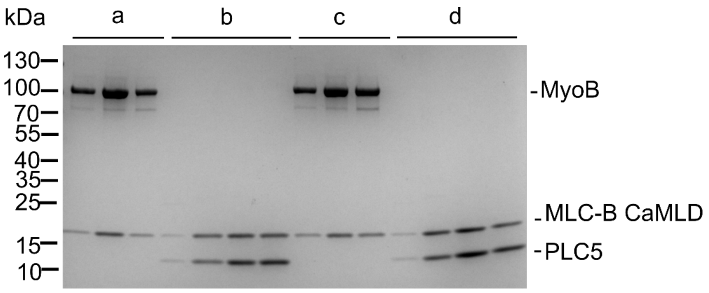


**Figure S3.** The incubation of MLC-B CaMLD and PLC5 with the MyoB-MLC-B CaMLD pre-formed complex. Elution fractions a containing MyoB-MLC-B CaMLD and elution fractions b containing MLC-B CaMLD and PLC5 (SEC buffer: 20 mM Tris pH 7.5, 150 mM NaCl, 1 mM $CaCl_2$, 5 mM $MgCl_2$, 0.5 mM TCEP). Elution fractions c containing MyoB-MLC-B CaMLD and elution fractions d containing MLC-B CaMLD and PLC5 (SEC buffer without $Ca^{2+}$).

### Tables

**Table S1:** The estimated SAXS invariants of MyoB-MLC-B CaMLD complex.

| Protein | MyoB-MLC-B CaMLD |
|---|---|
| $I_0$* (absolute, Guinier) | $3.33x10^3$ |
| $I_0$* (absolute, real space) | $3.25x10^3$ |
| $R_g$ (Å, Guinier) | 56.1 |
| $R_g$ (Å, real space) | 56.3 |
| $r_{avg}$ (Å) | 67.5 |
| $D_{max}$ (Å) | 208 |
| $V_{porod}$ ($nm^3$) | 285.3 |
| $P_x$ | 3.6 |
| MW (kDa), theoretical | 111.7 |
| MW (kDa), SAXSMoW # | 168.1 |
| DAMMIN $\chi^2$ | 1.3 |
| GASBOR $\chi^2$ | 1.3 |

**Table S2:** Secondary structure contents of MyoB-MLCB-CaMLD, MLC-B and Myosin V.

| Protein | α-helix (%) | β-sheet (%) | Turn (%) | Others (%) |
|---|---|---|---|---|
| Myosin B | 36 | 14 | 13 | 37 |
| MLC-B (8) | 40 | 14 | 17 | 27 |
| Myosin V (PDB: 1W7I, chain A) (20) | 41 | 14 | 11 | 34 |

**Table S3:** Melting temperature of MyoB-MLC-B CaMLD, Myosin-2 and β-cardiac myosin

| Protein | Conditions | $T_m$°C |
|---|---|---|
| MyoB-MLC-B CaMLD | nucleotide-free | 50.6 |
| *Dictyostelium discoideum* myosin-2 motor domain (54) | nucleotide-free | 45.6 |
| β-cardiac myosin motor domain (55) | nucleotide-free (low salt buffer) | 45.8 |
| | nucleotide-free (high salt buffer) | 46.5 |

### Movies

**Movie 1.** *In vitro* motility assay fluorescence microscopy of MyoB-MLC-B CaMLD (adsorption). Without the addition of ATP, the actin filaments bound to the surface and were non-motile. After ATP addition, the filaments started moving, showing ATP-driven motility.

**Movie 2.** *In vitro* motility assay fluorescence microscopy of MyoB-MLC-B CaMLD (antibody). Without the addition of ATP, the actin filaments bound to the surface and were non-motile. After ATP addition, the filaments started moving, showing ATP-driven motility.

**Movie 3.** *In vitro* motility assay fluorescence microscopy of MyoB without light chain (adsorption). Without the addition of ATP, the actin filaments bound to the surface and were non-motile. After ATP addition the filaments were immobile.
